# Apparent survival suggests the female-biased dispersal in a truly subterranean rodent, *Ellobius talpinus*

**DOI:** 10.64898/2026.07.30.741862

**Authors:** Anna E. Naumova, Andrey O. Fedosov, Varvara R. Nikonova, Arman M. Bergaliev, Margarita M. Dymskaya, Antonina V. Smorkatcheva

## Abstract

Adult sex ratios in mammals are usually female-biased, but the northern mole vole (*Ellobius talpinus*) (Pallas, 1770), a cooperatively breeding subterranean rodent, shows a persistent male bias that increases with age. Combining four years of capture-mark-recapture data with radiographic age estimation and genetic parentage data, we modeled apparent survival and state transitions (young-of-the-year, breeder, non-breeder) separately by sex. Young individuals showed significantly lower apparent survival than adults in both sexes, consistent with dispersal occurring predominantly in the first year of life. Within each age class, females showed consistently lower apparent survival than males, and this pattern held within breeding and non-breeding status groups alike, with female breeders showing lower apparent survival than male breeders. These results point to female-biased natal dispersal, an atypical pattern for mammals. We propose that faster turnover of reproductive females than males, in combination with singular breeding, favor female dispersal in search of vacant breeding positions, while promoting male philopatry and facultative polyandry.

## Introduction

In most natural populations, the adult sex ratio (ASR) tends to be biased (Donald, 2007; Székely et al., 2014; Pipoly et al., 2015) ranging from strongly male-biased to those composed exclusively of adult females (Székely et al., 2014; Cockburn et al., 1985). The bias in ASR may originate from multiple factors, including a skewed birth sex ratio (Miranda et al., 2025), sex-specific differences in growth or maturation rates (Crowley, 2000; Welbergen, 2010), or methodological artifacts arising from differential detectability or sampling probability between the sexes (Sandercock, 2006; Pickett et al., 2012; Romano et al., 2018). However, sex-specific survival rates appear to be the main cause of the ASR bias in most species. A wealth of accumulated evidence demonstrates distinct inter-taxon trends: males tend to be more numerous in bird populations (implying higher male survival), whereas females usually predominate in mammal populations. Several non-exclusive hypotheses have been proposed to explain the observed patterns (Székely et al., 2014; Payevsky, 2020; Staerk et al., 2025).

In a large-scale phylogenetic analysis of tetrapods, Pipoly et al. (2015) showed that ASR is consistently biased toward the homogametic sex, suggesting that this pattern may be driven by elevated mortality in the heterogametic sex (males in mammals and females in birds). The precise mechanisms remain unresolved and warrant further investigation. Additionally, this hypothesis does not account for the differential sex-biased mortality skew observed within each class (Pipoly et al., 2015).

Sex-biased adult survival may arise from asymmetry between males and females in reproductive investment, such as parental care or mating competition (Williams, 1966; Promislow, 1992; Liker and Székely, 2005; Froy et al., 2016; Romano et al., 2022; Staerk et al., 2025). Specifically, the sex that exhibits greater reproductive investment may experience elevated mortality as a consequence of physical exhaustion (Cockburn et al., 1985; Małek et al., 2023) or higher vulnerability to parasitism (Metcalf and Graham, 2018). Indeed, high mortality among reproductive females, resulting in a male-biased adult sex ratio, has been documented in many bird species (Romano et al., 2022 but see Winder et al., 2025). In contrast, studies on most mammals show that the turnover rate of reproductive females is usually lower than that of reproductive males (Promislow, 1992).

An additional, non-mutually exclusive explanation is that both the contrasting ASR biases typical of birds and mammals and the numerous deviations from these class-specific patterns reflect differences in natal dispersal strategies and the elevated mortality associated with the dispersing sex (males in mammals and females in birds) (Greenwood, 1980; Végvári et al., 2018; Payevsky, 2021). While sex-biased dispersal is often interpreted as a mechanism to avoid inbreeding, the evolutionary factors determining which sex disperses remain a subject of active debate (Perrin and Mazalov, 1999, 2000; Gros et al., 2009).

Differential survival and resulting skewed sex ratios create selective pressures that mold behaviors related to partner choice, mating competition, and parental investment (Donald, 2007; Grüebler and Naef-Daenzer, 2008; Liker et al., 2015; Székely et al., 2014), and there is recursive feedback between sex ratio, breeding systems and dispersal strategies (Perrin and Mazalov, 2000; Miranda et al., 2025). Therefore, data on variation in sex-specific survival across species are essential for understanding its ultimate and proximate causes and the consequences for the evolution of social systems. Species or populations with traits diverging from those of their taxonomic relatives warrant particular scientific attention. Such comparative studies enable more precise identification of the predictors underlying ecological patterns.

The northern mole vole (*Ellobius talpinus*) (Pallas, 1770) is a truly subterranean rodent, inhabiting the grasslands of the Palearctic (Kryštufek and Shenbrot, 2022). Northern mole voles from two relatively well-studied populations (Ural region - Evdokimov, 2001, 2013; Evdokimov and Pozmogova, 1992; Shevlyuk and Yelina, 2008; Low Volga region - Bergaliev et al., 2026) exhibit unusual demographic and behavioral traits that make them unique within the subfamily *Arvicolinae*. First, northern mole voles live in cooperative groups of up to >25 individuals, with reproduction usually limited to one female and one (sometimes two or three) male. Second, its longevity far exceeds typical arvicoline standards, reaching over 6 years in mole voles (Evdokimov, 1997, 2001). Third, males predominate in these populations, with their proportion increasing in progressively older age classes (Evdokimov, 1997; Evdokimov and Pozmogova, 1984, 1992, 1993; Bergaliev et al., 2026), defying the typical mammalian pattern seen in most voles (Jannett, 1981; Myers and Krebs, 1971; Briner et al., 2007) and subterranean rodents from other taxa (Zenuto, 1998; Busch et al., 2000; Šumbera et al., 2007; but see Brett, 1991; Finn et al., 2022).

Sufficient data to elucidate the mechanisms underlying this sexual structure remain unavailable. First, maintaining energy balance poses a significant challenge for breeding female subterranean rodents, as they must allocate limited resources to the competing demands of reproduction and digging (Vleck, 1979; Zenuto, 1998; Novikov, 2007). As a result, reproductive females may suffer from higher mortality than same-age males. However, in cooperative breeding systems, males and non-reproductive females may take on part of the energetic burden of the breeding female. This reduction in workload can lower the survival costs of reproduction, potentially leading to more equal mortality rates (Francioli et al., 2020; Houslay et al., 2020).

Another possible source of the sex-specific mortality and primary driver of the male-biased sex ratio in mole vole populations are a female-biased natal dispersal (Evdokimov, 2001). Most subterranean rodents are known or supposed to disperse above ground (Hazell et al., 2000; Finn, 2021), and this appears to be the case for mole voles as well (Evdokimov, 2013). Moreover, genetic data suggest the potential for long-distance dispersal (Rudyk et al., 2025), which is undoubtedly possible only above ground and must be associated with a very high risk.

It is not currently possible to assess, based on published data, the relative contribution of two potential sources of sex-specific mortality - sex-biased dispersal and reproductive costs - to the maintenance of the demographic structure documented for *E. talpinus*. Should female-biased dispersal be confirmed in this species, a thorough investigation of the underlying ecological, demographic, and social factors would be warranted to elucidate both the drivers and consequences of this mammalian atypical strategy, ultimately contributing to a broader understanding of how dispersal strategies evolve.

A population of northern mole voles from the Lower Volga region was subjected to long-term monitoring via capture-mark-recapture, non-invasive age estimation, and genetic parentage analysis. In this article, we analyze the population’s sex and age structure and apparent survival to address the following questions: (i) whether reproductive females suffer higher mortality than non-reproductive adults, thereby driving a male-biased adult sex ratio; and (ii) whether the sex difference in non-breeder apparent survival is consistent with female-biased natal dispersal.

## Materials and Methods

### Study area and sampling

Field work was conducted in the Saratov Region (Russia), four kilometers west of Dyakovka village (50.71°N, 46.71°E), on a study area of approximately five hectares. This territory lies on the border of steppe and semidesert zones. The vegetation cover is represented by the psammophytic-steppe and meadow-steppe types of plant communities (Vasukov et al., 2018).

In the Lower Volga region, mole vole reproduction follows a seasonal pattern, with parturition occurring from late winter/early spring through mid-summer. Both male and female mole voles seem to delay reproduction until after their first winter (Evdokimov, 2001; Shevlyuk and Yelina, 2008; Bergaliev et al., 2026). Each breeding season, females can bear up to three litters, with litter size ranging from 2 to 8 offspring (Shevlyuk and Yelina, 2008 for the South Ural population). Throughout the study period, local population density remained high, ranging approximately 15–30 individuals/ha.

Capture-mark-recapture surveys were conducted from July 2021 to September 2025, with 2 – 4 trapping sessions carried out annually, typically from April to September in years following the initial survey. Each session ranged from 6 to 25 days in duration, with a mean of 14 ± 7 days.

The animals were trapped with metallic-spiral live-traps (Golov, 1954) with modifications) baited with carrot. The traps were checked each 15-20 minutes. For each captured individual, the following data were recorded: capture date and time, GPS coordinates of the capture location (precision to the nearest 0.0001 decimal degree), sex, pelage condition (grayish, brown or undergoing molt), and body mass (to the nearest 0.1 g). The combined width of the upper incisors was measured using a digital caliper to the nearest 0.01 mm and employed to distinguish juveniles from adults (Kuprina and Smorkatcheva, 2019 for a closely related species, *Ellobius tancrei* (Blasius, 1884)).

At first capture, each animal was tagged with a 1.25*7 mm microtransponder (Star Security Technologies Co., Shanghai, China). In addition, distal phalanges from one or two toes were collected, immediately preserved in 96% ethanol, and stored at −20°C for subsequent analysis of the population genetic structure and of parentage (Rudyk et al., 2025; Bergaliev et al., 2026).

Since July 2022, radiography of animal skulls with portable X-ray equipment (Rexstar LCD, Korea) and the Dental Radiovisiography Sensor EzSensor 1.5 (Vatech) has been added to the above manipulations. The radiographs were later used to categorize the animals into age classes (see below and Nikonova et al., 2024 for details). In addition, bioacoustic data were collected during some trapping sessions (Dymskaya et al., 2025).

From 2023 to the present, a commercial microtransponder scanner (CMS) designed for livestock applications (PR-250S, PARTNER VET, Saint Petersburg, Russia) was used. The CMS is a scanning probe that is inserted into the burrow and requires constant operator supervision. Since 2024, we have used a custom-built reader (CiberGato) designed to automatically record the ID of the tagged animals moving through tunnels (Fedosov in prep.).

Since July 2021, a total of 467 individuals have been marked in the study area.

### Age class determination

Animals with combined width of the upper incisors ≤ 3.15 mm and/or those with grayish fur or showing signs of juvenile molt were identified as young-of-the-year (Nikonova et al., 2024). Ages of animals with joint width of the upper incisors > 3.15 caught in 2022– 2024 were further assessed based on their radiographs. The analysis procedure is briefly described below (see Nikonova et al., 2024 for details).

A total of 578 images from 377 individual animals were stored and processed using EzDent i v.3.0.8.0 Console software (EWOOSOFT Co., Ltd., Korea http://ewoosoft.com). On the X-ray images, the lengths of the second synclinal fold of the first upper molar and that of the first lower molar were measured (accuracy of 0.01 mm) with a digital tool (Supplementary Fig S1; Nikonova et al., 2024). Next, a principal component analysis (PCA) was performed to combine two highly correlated metrics into a new variable (molar condition, MC). Subsequently, discriminant function analysis (DFA, *lda* function from the MASS package in R4.4.3 -Venables and Ripley (2002)) was conducted to classify all images into three age classes: class 1 - images from young-of-the-year; class 2 - images from yearling; class 3 – images from animals who survived at least two winters. MC and the Julian date of the X-ray were used as predictors. A total of 236 *known-class* radiographs from 127 animals (class 1: n = 145; class 2: n = 38; class 3: n = 53) were used as the training set. Prior probabilities were based on group size. The test error was estimated using leave-one-out cross-validation (*caret* package in R 4.4.3). Finally, we applied the obtained classification functions to classify 342 *unknown-class* images from 263 individuals. For the purposes of this study, all overwintered animals (classes 2 and 3) were categorized as adults.

### Breeding status determination

In the field, females were recorded as reproductive if they displayed an enlarged abdomen (suggestive of pregnancy) and/or enlarged teats (indicating lactation). In 2022 - 2024, female visual examination data were supplemented with genetic data (Bergaliev et al., 2026). Male reproductive status could not be assessed externally (Shevlyuk and Yelina, 2008; our observations); thus, it was based solely on parentage analysis (2022 - 2024). The reproductive status of males recorded in 2025 remains unknown.

The parentage analysis methods are described in detail in Bergaliev et al. (2026). Shortly, we genotyped 334 individuals (91% of the animals sampled in 2021 - 2024) using a panel of 10 previously developed microsatellite loci. Then, the full-pedigree likelihood method implemented in the software COLONY v. 2.0.7.1 (Jones and Wang, 2010) was used for the parentage and sibship assignment. These analyses were conducted separately for each study year. The reliability of the obtained relationships was evaluated (i) by comparing the assigned mother-offspring and sibling pairs with those expected from trapping records and (ii) by mitochondrial D-loop haplotype matching.

### Statistical analysis

Analyses were restricted to 2022 - 2025 data, since age was known for only a small proportion of individuals captured in 2021. Of the 428 northern mole voles captured over the four-year study period, a total of 140 individuals were excluded for the following reasons: unknown age class at any study period (n = 40), unknown sex (n = 3). From the analysis of apparent survival rates, we also excluded animals that were exclusively caught during the last year of the study (2025), precluding any contribution to the analysis (n = 93), and animals that died during capture from overheating or stress (n = 4).

The inclusion of transmigrants (individuals temporarily present in the population without permanent settlement) could bias apparent survival estimates downward by inflating perceived mortality through temporary disappearance rather than true death or permanent emigration. Combination of trapping and genetic kinship data revealed that throughout 2022 – 2024 there were no transmigrants.

Two-tailed binomial tests (R v. 4.3.1; R Core Team 2026) were used to assess deviations from an equal sex ratio separately for each age class (young vs. adults) and breeding status (breeders vs. non-breeders, adult animals only) within each study year (2022 - 2025). For breeders, however, tests were restricted to 2022 - 2024, as the number of reproductive males in 2025 was unknown.

To address the questions outlined in the Introduction, multistate mark-recapture model set (Lebreton and Pradel, 2002) was constructed using RMark package (v. 3.0.7; Laake, 2013) in R (v. 4.5.2; R Core Team 2026), and run in program MARK (White and Burnham, 1999). The fundamental input to multistate models is the encounter histories (series of symbols where the letters indicate an encounter of an animal in a specific state on a given sampling occasion and zeroes indicate non-detection of the animal; e.g., CBB0).

As parameters of interest were estimated over annual time intervals and state of the animal was constant during this interval, multiple encounters of the same animal within the same sampling season (May - September) were pooled into a single occasion. These are probabilities of apparent survival (*φ*), encounter (p) and transition (*ψ*). Apparent survival probability reflects true survival reduced by permanent emigration, as individuals that permanently leave the study area cannot be distinguished from those that have died. Encounter probability reflects the probability of detecting a living marked individual on a given occasion, which was fixed to 1 as all marked individuals were detected at each annual occasion or had permanently left the population. Transition probability describes the probability of an individual moving from one state to another between consecutive occasions, conditional on survival.

Four mutually exclusive states were defined based on age and reproductive status: young-of-the-year (Y), reproductive (B), non-reproductive (N) and unknown (U). Possible transitions included young-of-the-year to reproductive, young-of-the-year to non-reproductive, reproductive to reproductive, reproductive to non-reproductive, non-reproductive to reproductive and non-reproductive to non-reproductive. Unknown status was only applied in the final sampling occasion (2025) to all males (animals from each state transitioned to U, no transitions from U were modelled). Transition from reproductive to non-reproductive was not observed in females and its probability was therefore set to zero.

To estimate sex, age and reproductive status effect on apparent survival probability (*φ*) and transition probability (*ψ*), a global model structured as *φ* (status * sex + time) *ψ* (status: tostatus: sexCtime) p (1) and its nested models were constructed. Sex was added as a group variable. Time was included as a nuisance parameter to account for annual variation in survival not attributable to state or sex. sexCtime was a covariate combining sex and time, allowing transition probabilities to be estimated independently for each sex while distinguishing the final sampling occasion from all preceding ones for male animals. Model notation follows Lebreton et al. (1992), with ‘*’ denoting an interaction plus additive effect, ‘+’ an additive effect and ‘:’ an interaction only effect.

To estimate the overdispersion factor in the case of non-systematic deviations (‘extra-binomial variation’), we used the median ĉ approach implemented in the program MARK (White and Burnham, 1999) for the global model. No evidence of overdispersion was found (ĉ = 0.97) and model selection was therefore conducted without adjustment for overdispersion.

Model selection was based on Akaike’s information criterion adjusted for small sample size (AICc; Burnham and Anderson, 2002). The Akaike weight of the top model was below 0.9, suggesting considerable model selection uncertainty. Therefore, we used function model.average in RMark to obtain averaged parameter estimates and standard errors unconditional on a given model (Buckland et al., 1997; Burnham and Anderson, 2004).

## Results

### Age class determination

PCA yielded PC1 which accounted for 96% of the total variation (factor loading 0.98). In the following, we used PC1 as an indicator of molar condition (MC).

DFA based on 236 known-class radiographs confirmed that the molar condition and day of radiography, taken together, ensured discrimination between age classes (Wilks’ lambda = 0.12; χ2 = 215.9; df = 4; p < 0.0001). The plot of the canonical scores for the first two discriminant functions illustrates the separation among age classes (Fig. 1). The first discriminant function was highly correlated (r = 0.89) with the molar condition, whereas the second discriminant function was highly correlated with the day of radiography (r = 0.99). The first discriminant function accounted for 99.9% of the grouping variation.

**Fig. 1.**
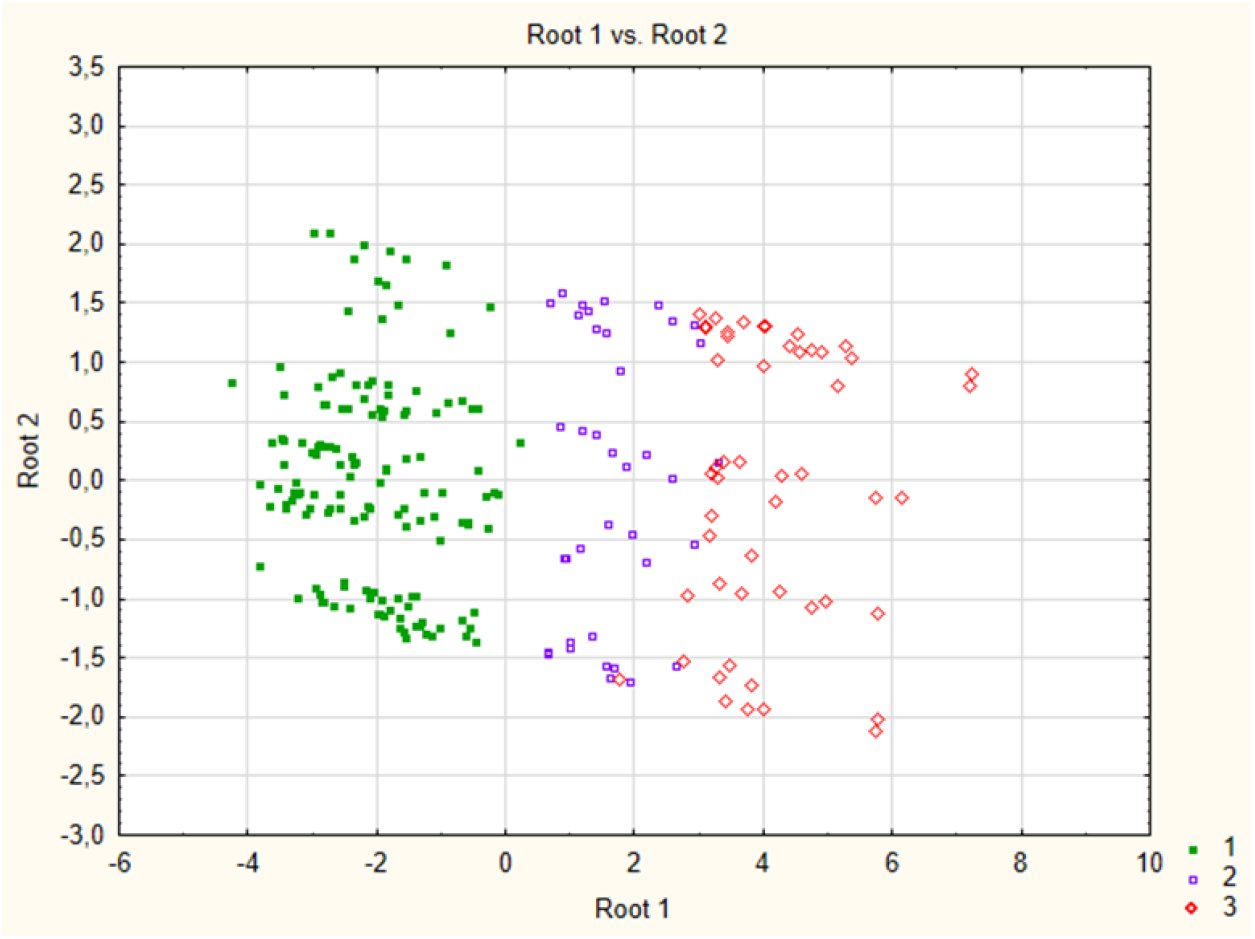
Scatterplot illustrating DFA results for classifying 236 known-class radiographs. 1 - 3 – known age classes of the radiographs. The graph was produced using Statistica 12.

The following discriminant equations were obtained:

Class 1: -6.28 + 7.03 molar condition + 0.06 day.

Class 2: -3.47 - 2.47 molar condition + 0.02 day.

Class 3: -8.57 - 9.14 molar condition - 0.00007 day.

The predictive accuracy of the model for the analysis sample was 0.97. The predictive accuracy of the cross-validation sample also was 0.97 (95% CI: 0.943 - 0.989).

Only one radiograph from class 1 was mistakenly assigned to class 2, four images from class 2 were mistakenly assigned to class 3, and two images from class 3 were mistakenly classified as class 2. Three out of the seven incorrectly classified images belonged to the same individual. All but two misclassifications had low posterior probabilities (<0.7), therefore age assignment that had posterior probabilities >0.80 was considered reliable. Using the obtained discriminant function, we were able to identify the age class for 292 images from 239 individuals and to exclude one of two age classes for the remaining 50 images from 45 individuals. All age estimates for repeatedly radiographed individuals were consistent across images: the images from the same individual were always assigned to the same class if they had been obtained in the same year and to different classes if they had been obtained in different years. As a result, among the animals captured in the focal area, 70 individuals in 2022, 126 in 2023, 168 in 2024, and 143 in 2025 could be unambiguously classified as either a young-of-the-year or an adult (Table 1), 387 individuals in total.

**Table 1.** The number of animals of each sex in each state each year: F – female, M – male, Y – young-of-the-year, B – breeder (% of all adults of the same sex in parentheses), N – non-breeder, U - unknown. REM column provides the number of animals removed from the sample on specific year (% of total number in parentheses).

|  | F Y | F B | F N | M Y | M B | M N | M U | REM |
| --- | --- | --- | --- | --- | --- | --- | --- | --- |
| 2022 | 22 | 6 (75) | 2 | 25 | 7 (50) | 7 | - | 20 (22) |
| 2023 | 36 | 8 (38) | 13 | 41 | 6 (23) | 20 | - | 11 (8) |
| 2024 | 44 | 13 (68) | 6 | 57 | 15 (33) | 30 | - | 9 (5) |
| 2025 | 40 | 7 (35) | 13 | 31 | - | - | 51 | 7 (5) |

**Table 2.** Candidate model set to estimate apparent survival (*φ*) and transition probabilities (*ψ*) of mole voles captured between 2022 and 2025. The table includes the model, number of model parameters (Npar), AICc, delta AICc (ΔAICc), model weights (Weight) and model deviance (Deviance).

| Model | Npar | AICc | ΔAICc | Weight | Deviance |
| --- | --- | --- | --- | --- | --- |
| $\varphi$ (~status + sex)<br>$\psi$ (~status:tostatus:sexCtime) | 9 | 512.94 | 0.00 | 0.596 | 60.68 |
| $\varphi$ (~status + sex + time)<br>$\psi$ (~status : tostatus : sexCtime) | 11 | 514.99 | 2.05 | 0.214 | 58.48 |
| $\varphi$ (~status * sex)<br>$\psi$ (~status : tostatus : sexCtime) | 11 | 516.83 | 3.89 | 0.085 | 60.32 |
| $\varphi$ (~sex) $\psi$ (~status : tostatus : sexCtime) | 7 | 518.00 | 5.06 | 0.047 | 69.94 |
| $\varphi$ (~status * sex + time)<br>$\psi$ (~status : tostatus : sexCtime) | 13 | 518.96 | 6.03 | 0.029 | 58.15 |
| $\varphi$ (~sex + time)<br>$\psi$ (~status : tostatus : sexCtime) | 9 | 520.51 | 7.57 | 0.014 | 68.25 |
| $\varphi$ (~status)<br>$\psi$ (~status : tostatus : sexCtime) | 8 | 520.98 | 8.04 | 0.011 | 70.83 |
| $\varphi$ (~status + time)<br>$\psi$ (~status : tostatus : sexCtime) | 10 | 523.25 | 10.32 | 0.003 | 68.88 |
| $\varphi$ (~1) $\psi$ (~status : tostatus : sexCtime) | 6 | 528.12 | 15.18 | 0.000 | 82.14 |
| $\varphi$ (~time) $\psi$ (~status : tostatus : sexCtime) | 8 | 530.93 | 17.99 | 0.000 | 80.78 |

### Population sex ratio and the breeder proportions

Demographic structure of the focal population is summarized in Table 1. The sex ratio among young-of-the-year did not differ from 1:1 in any of the study years with a slight deviation toward males in 2022–2024 and females in 2025 (2022: p = 0.763, 2023: p = 0.654, 2024: p = 0.271 and 2025: p = 0.400). The sex ratio among adult animals was consistently biased towards males, although significant deviation from 1:1 sex ratio was revealed only in 2 study years (2022: p = 0.191; 2023: p = 0.375; 2024: p = 0.002; 2025: p = 0.0003) (Table 1, Supplementary Fig. S3).

The proportion of breeders varied across years and between sexes. The number of breeding males and females was comparable across the three years available for comparison (2022 - 2024), and none of the tests revealed a statistically significant deviation from an equal (1:1) sex ratio among breeders (2022: p = 0.500; 2023: p = 0.790; 2024: p = 0.702) (Table 1).

### Apparent survival

Among the candidate model set, the top-ranked model *φ* (sex + status) *ψ* (status: tostatus: sexCtime) received substantial support (weight = 0.596). The summarized weight of models containing sex and stratum as additive effects on apparent survival was 0.925, indicating strong and consistent support for both variables. Notably, time was not retained in the top model, suggesting that annual variation contributed little to explaining apparent survival in this population beyond previously mentioned effects, although it cannot be fully excluded (Burnham and Anderson, 2002). Detailed numerical annual survival rate, transition probabilities and 95% confidence intervals for each demographic group are provided in Supplementary Tables S1, S2. Estimates of apparent survival and transition probabilities combined for the 2023 - 2024 time period is illustrated on Fig. 2 A, B. Combined estimates for other periods are provided on the Supplementary Fig. S2 A, B, C.

**Fig. 2.**
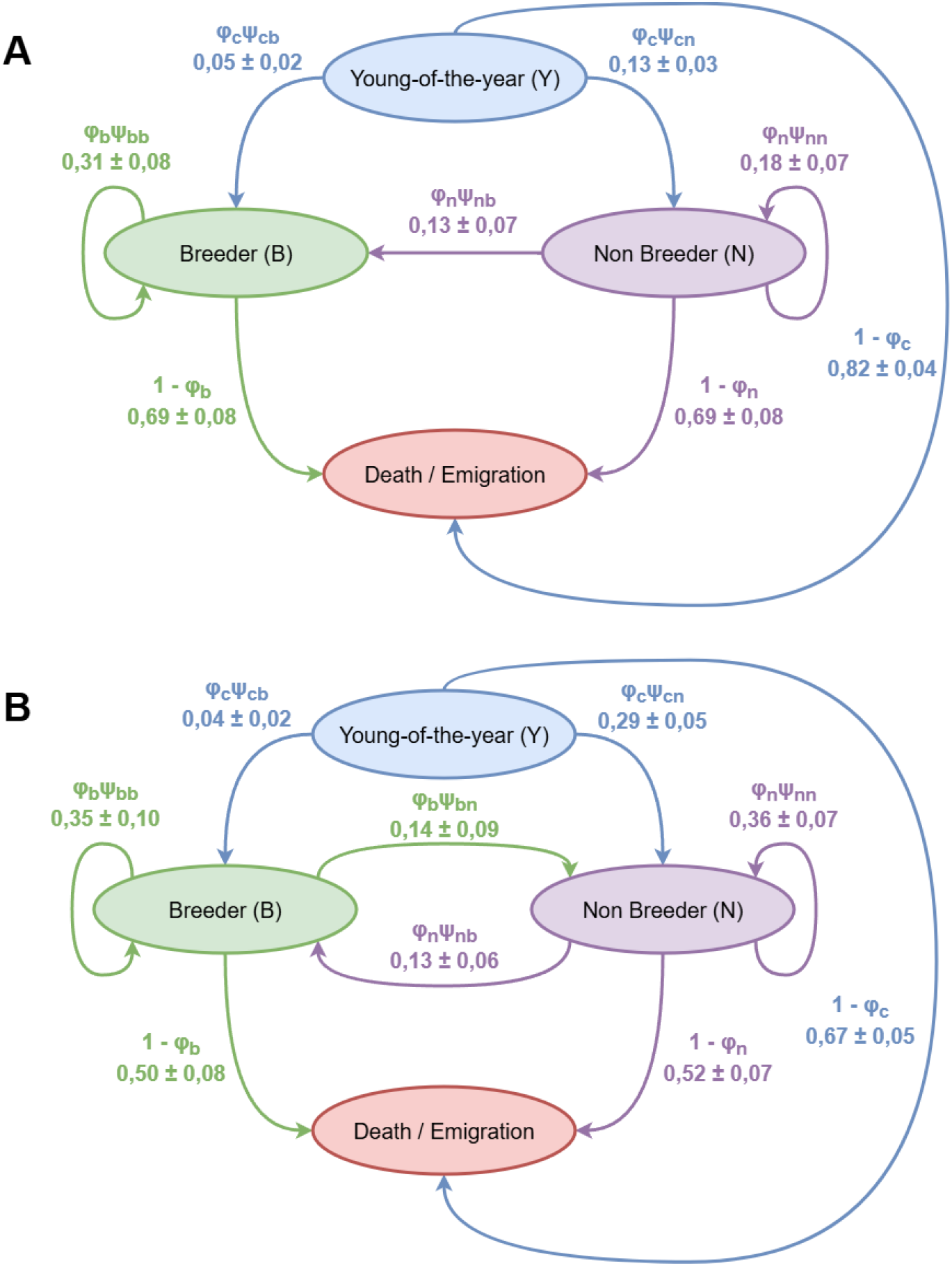
**A, B** Graphical representation of apparent survival and transition probabilities combined for A: females (n = 128) and B: males (n = 160) in time period between 2023 and 2024 based on model averaging, numbers are settled as value ± SE.

Females showed substantially lower apparent survival than males. This difference was most pronounced in young-of-the-year, with female apparent survival being at nearly half the rate of male (*φ*_c_ = 0.18 ± 0.04 and *φ*_c_ = 0.33 ± 0.05 respectively for 2023 - 2024 time period). Among adults, breeder and non-breeder apparent survival was similar within each sex (*φ*_b_ = 0.31 ± 0.08 and *φ*_n_ = 0.31 ± 0.08 for females and *φ*_b_ = 0.50 ± 0.09 and *φ*_n_ = 0.48 ± 0.08 for males for 2023 - 2024 time period).

Female breeders showed complete state fidelity - no transitions to non-breeder state were observed - while male breeders showed variation (*ψ*_bb_ = 0.71 ± 0.17).

Young-of-the-year of both sexes predominantly transitioned to non-breeder status, though this tendency was stronger in males (*ψ*_cn_ = 0.88 ± 0.07) than females (*ψ*_cn_ = 0.71 ± 0.11). Adult non-breeders tended to remain in their current state in both sexes, with males showing stronger state fidelity (*ψ*_nn_ = 0.73 ± 0.11) than females (*ψ*_nn_ = 0.57 ± 0.19).

## Discussion

We provide the first apparent survival estimates ever reported for any *Ellobius* species, along with the first demographic structure analysis for this species in the Lower Volga Region. To assess effects of age, sex, year, and reproductive status on apparent survival rate, we employed capture-mark-recapture methods, minimally invasive age estimation, and genetic data. Despite an approximately equal sex ratio among juveniles, adult males outnumbered adult females in our population, suggesting higher survival in males compared to females. Our results align with previous studies reporting a slight male predominance in most northern mole vole populations (Orenburg Region - Yelina, 2006; Central Kazakhstan - Shubin, 1961; Bashkortostan, Chelyabinsk Region, Northern Kazakhstan - Evdokimov and Pomozgova, 1984; Evdokimov, 2013; Altai Krai – Novikov et al., 2007 but see Novikov et al., 2007 for the population from the Novosibirsk Region). The male-biased sex ratio is consistent with the results of our apparent survival analysis, which showed lower apparent survival in females than in males within the same age category. Given that apparent survival does not differentiate between mortality and permanent emigration (Lebreton et al., 1992), both processes warrant consideration.

Predator disturbance of burrows and nests, and abiotic stressors such as flooding, frost, and ice-crusted ground, are not sex-specific and therefore cannot account for the systematically lower survival of non-reproductive females relative to males. Among the potential causes of elevated mortality in females several key factors can be identified. First, intrasexual competition among females for reproductive status and social suppression of subordinate individuals by the dominant breeding female may result in chronic stress and, consequently, increased mortality (Sharp and Clutton-Brock, 2011; Maag et al., 2022) Indirect support for this mechanism is provided by laboratory data on a closely related species, the Zaisan mole vole (*Ellobius tancrei*), in which elevated levels of female competition have been documented. However, the authors suggest that under natural conditions this would most likely manifest as emigration of subordinate individuals rather than their death (Smorkatcheva and Kuprina, 2018).

A second potential source of increased female mortality is a higher level of burrowing activity among non-reproductive females relative to males, entailing greater energetic expenditure and possibly increased exposure to adverse environmental conditions. However, this explanation appears unlikely. Data from social bathyergids, a rodent group with similar ecological specialization and social system, suggest that sex differences in cooperative burrowing activity among non-breeders are largely absent across species (Zöttl et al., 2022). If a similar pattern holds for mole voles, differential energetic expenditure associated with burrowing is unlikely to account for the observed sex difference in apparent survival among adults. However, investigating the activity patterns of different strata within our focal *Ellobius talpinus* populations represents the next planned step in our research.

Two most parsimonious explanations for the observed sex difference in apparent survival are high reproductive costs in females, female-biased dispersal, or a combination of both factors. The tradeoff between female reproduction and survival has been documented in several species of mammals (Speakman, 2008; Fisher and Blomberg, 2011; McHuron et al., 2023; Ginther et al., 2024). Dispersal is likewise commonly associated with high mortality rates; in small mammals, the survival of dispersing individuals is nearly 50% lower than that of philopatric individuals (Johnson and Gaines, 1990).

If reproductive costs were the primary driver, breeding females would be expected to show lower apparent survival than non-breeding females and breeding males. Our data did not support the first prediction: apparent survival estimates for adult breeder and non-breeder females were essentially identical. At the same time, breeding males’ apparent survival rate was significantly higher than this of breeding females (0.50 vs. 0.31), although the probability of retaining reproductive status did not differ between the sexes. Unlike females, males that lost breeder status sometimes survived as non-breeders; in one such case, breeding status was subsequently inherited by the male’s sons (Bergaliev et al., 2026).

In non-reproductive animals, comparisons across sexes revealed the same clear pattern: females had lower apparent survival than males, with young-of-year females showing the lowest apparent survival of all. These findings are consistent with the hypothesis that sex-biased dispersal contribute to the observed sex-biased mortality.

The sex difference in dispersal level may be expressed as unequal disperser fractions (e.g. Graw et al., 2016), unequal dispersal age (e.g. Puzachenko, 1998) and/or unequal dispersal distance (e.g. Mabry et al., 2013). According to Evdokimov (1999), northern mole voles from the Ural population show no pronounced sex bias in overall dispersal rates, but females dispersed predominantly at younger ages (young and yearlings), whereas males tended to disperse at older ages (two-year-olds). In our population, adults of both sexes show higher apparent survival rates than young-of-the-year. This suggests that dispersal in both sexes occurs predominantly during the first year of life, albeit evidently at different rates. The apparent survival estimates obtained for both age categories of non-reproductive animals suggest that the proportion of emigrants is higher among females than among males in both the first and subsequent years of life. Our genetic data for the study population (Bergaliev et al., 2026) indicate that philopatric reproduction is a very rare phenomenon in both sexes. Over four years of monitoring, only 7 cases were recorded for males and 3 for females which appears to be consistent with male-biased philopatry.

The dispersal costs and risks predictably increase with the distance travelled (Johnson and Gaines, 1990). In some mammalian species, despite an equal dispersal sex ratio, one sex consistently disperses over greater distances than the other (Roy et al., 2014). As a consequence, the ratio of successful immigrants tends to be biased toward the sex dispersing over shorter distances. Studies on tuco-tucos (Cutrera et al., 2005) and surface-dwelling voles (Davis-Born and Wolff, 2000; Gauffre et al., 2009) have demonstrated that males and females frequently differ in the distances they cover during dispersal, implying that the two sexes are exposed to different levels of dispersal-associated mortality risk. Long-term studies conducted on the Ural population of the northern mole vole have previously shown that males exhibit a preference for local movements compared to females (Evdokimov, 1999). Our data from the focal site are insufficient to test sex-specific differences in dispersal distance. Over the four years of study, we reliably documented only three inter-family dispersal events within the 5-ha focal plot (2 males and one female, see Bergaliev et al., 2026, Supplementary Table S4); given this negligible sample size, no inference can be drawn regarding the direction or magnitude of sex bias in dispersal distance. Similarly, no long-distance emigrants were detected beyond 100 m from the focal site using an automated microtransponder readers, which may reflect limited number of placements rather than a true absence of long-distance dispersal.

Kinship analysis showed that among animals born during our period of intensive trapping, only a few individuals lacked close relatives (parents, siblings or half-siblings) and could thus be considered potential immigrants (Bergaliev et al., 2026, Supplementary Tables S3 and S4). Moreover, most of these were captured at the periphery of the focal area, suggesting that their parents may have evaded trapping. Only 7 females and 1 male could be reliably considered as immigrants from outside our focal area. The extremely low number of immigrants, together with the rarity of intra-site dispersal and low apparent survival rates, points to very high mortality during dispersal, suggesting that emigrants of both sexes likely undertake long-distance dispersal.

The previous genetic studies on the focal population found no obvious sex-specific pattern in the spatial distribution of mitochondrial haplotypes, providing no clear evidence of sex-biased dispersal. (Rudyk et al., 2025). This discrepancy may reflect methodological differences: whereas Rudyk et al. (2025) focused on immigrants that contributed genetically to the population, the present study addresses emigrants - animals that departed from the population. These two quantities are not necessarily equivalent, as numerous examples from subterranean rodents demonstrate that one sex may disperse over shorter distances, thereby achieving greater immigration success. It is also possible that the absence of concordance between these findings is partly attributable to limited sample sizes. Notably, for the Saratov population, the authors likewise proposed long-distance dispersal.

Female-biased dispersal is highly atypical for mammals (Greenwood, 1980, Mabry et al., 2013). Male-biased dispersal is the predominant strategy in both surface-dwelling voles and subterranean rodents, regardless of social organization (Boonstra et al., 1987; Getz et al., 1994; Lambin, 1994; Salvioni and Lidicker, 1995; Сutrera et al., 2005; Fernández-Stolz et al., 2007; Borkowska, 2010; Mora et al., 2010; Patzenhauerová et al., 2010; Le Galliard et al., 2012; Bray et al., 2012). This pattern in voles can be modified by ecological context. Sex bias in dispersal can weaken or disappear under island isolation and high density (beach vole, *Microtus breweri* (Baird, 1858) (Tamarin, 1977)) or resource saturation (California vole, *M. californicus* (Peale, 1848) (Heske, 1987)). It can even reverse for specific dispersal types or life stages, as shown in the common vole, *M. arvalis* (Pallas, 1778), in which short-distance dispersal is male-biased, whereas long-distance dispersal is sex-balanced (Gauffre et al., 2009).

Among social subterranean rodents, male-biased dispersal also seems to predominate. It has been reported in the social tuco-tuco, *Ctenomys sociabilis* (Pearson and Christie, 1985) (Lacey and Wieczorek, 2004), common mole-rat, *Cryptomys natalensis* (Lesson, 1826) (Hazell et al., 2000; Finn, 2022), naked mole rat*, Heterocephallus glaber* (Rüppell, 1842) (O’Riain et al, 1996) and Mechow’s mole-rat, *Fukomys mechowii* (Peters, 1881) (Kawalika and Burda, 2007). However, some species show roughly equal dispersal between the sexes, for example, in the Damaraland mole-rats, *Fukomys damarensis* (Ogilby, 1838) (Finn et al., 2022), but males may disperse over greater distances than females (Mynhardt et al., 2021).

To our knowledge, female dispersal has not been reported in any social subterranean rodents. In a solitary subterranean rodent, the blind mole-rat, *Spalax microphthalmus* (Gueldenstaedt, 1770), dispersal bias is age-dependent: female-biased among yearlings, while males disperse predominantly in the second year and older age cohorts (Puzachenko, 1998). Against this background, the female-biased dispersal inferred for the *Ellobius talpinus* is unusual and requires explanation.

Three main non-mutually exclusive hypotheses have been proposed to explain the evolutionary causes of sex-biased dispersal patterns. The inbreeding avoidance hypothesis predicts that the sex experiencing greater costs of inbreeding depression - typically the sex investing more in offspring - should be the dispersing sex (Waser et al., 1986). The dual and contradictory nature of these predictions makes this hypothesis inherently difficult to test (Lehmann and Perrin, 2003). The resource competition hypothesis posits that the sex whose reproductive success is more strongly dependent on familiarity with and access to high-quality resources (such as food and shelter) will tend to be philopatric, while the sex whose success is primarily determined by the number of mating partners will be more likely to disperse (Clark, 1978; Greenwood, 1980). In mammals, females compete for food and shelter, so familiarity with the natal territory is beneficial, and females tend to be more territorial than males. Males, in contrast, compete primarily for mates and benefit from dispersing in search of new partners. (Clark, 1978; Greenwood, 1980; Mabry et al., 2013). Consequently, this hypothesis fails to account for the female dispersal documented in some polygynous and promiscuous mammals (e.g. hamadryas baboon, *Papio hamadryas* (Linnaeus, 1758) (Hammond et al., 2006), North American porcupine, *Erethizon dorsatum* (Linnaeus, 1758) (Sweitzer et al., 1998)). In *E. talpinus*, a subterranean species with facultative social and genetic polyandry, the reproductive success of the reproductive female may depend on the number of partners if they contribute substantially to the construction and maintenance of foraging tunnels. In support of this hypothesis, laboratory observations have shown that males spend more time digging than reproductive females in the closely-related *E. tancrei* (Smorkatcheva and Kumaitova, 2013), as well as in the subterranean (though morphologically weakly specialized) mandarin vole, *Lasiopodomys mandarinus* (A. Milne-Edwards, 1871) (Smorkatcheva, 2003). Long-term studies combining monitoring of reproductive groups with genetic analysis could help clarify the fitness consequences of polyandry for mole vole reproductive females.

The intrasexual competition hypothesis proposes that the dispersing sex is the one experiencing higher levels of within-sex competition for reproductive opportunities (Brom et al., 2016). Male-biased dispersal has been reported for several polygynous or promiscuous *Ctenomys* species (Lacey and Wieczorek, 2004; Cutrera et al., 2005; Mora et al., 2010) and *Heliophobius argenteocinereus* (Peters, 1846) (Patzenhauerová et al., 2010). The intrasexual competition hypothesis also plausibly applies to our focal population, where only a single female breeds per family while several males may share reproduction with unequal contributions (Bergaliev et al., 2026). Records of within-group polyandry, unusual traits of male anatomy and physiology as well as the slight female-biased size dimorphism suggests stronger female-female than male-male competition for breeding positions and predicts female-biased dispersal, consistent with our findings.

Although data on the northern mole voles support a link between female dispersal and high female–female competition, as well as with social or genetic polyandry, the direction of this relationship remains unclear. Despite the striking similarity reproductive systems and social structure between mole voles and social mole-rats (Bergaliev et al., 2026), the sex-specific strategies of these rodents differ. In *Fukomys* and *Cryptomys*, males are the competing and dispersing sex (Kawalika and Burda, 2007; Finn, 2022). We speculate that the key feature of the focal mole vole population driving this partial reversal of sex roles is a higher turnover of queens compared to reproductive males. Indeed, reproductive female disappearance appears to be a key trigger for both daughter dispersal and the onset of breeding among philopatric sons (Bergaliev et al., 2026), supporting the idea that female reproductive tenure is a central determinant of these sex-specific dispersal strategies. This contrasts with the situation in the common mole-rat, *Cryptomys natalensis* (Finn, 2022), and *F. damarensis* (Patzenhauerová et al., 2013) where breeding-female turnover is considerably lower than that of breeding males. In the present study, we did not detect sex differences in apparent survival among breeders that retained their breeder status; however, our analysis included all males that sired at least one offspring, i.e., both dominant and subordinate males from polyandrous groups (Bergaliev et al., 2026). Dominant males appear to survive and retain their status better than queens do. A dispersing female therefore has a higher probability than a dispersing male of encountering a strange family with an available breeding vacancy. On the other hand, a philopatric daughter whose mother has died faces either confrontation with an immigrant female or, at best, the opportunity for inbred reproduction with her father; both options are clearly worse than the prospects of a disperser. For males, then, the situation is reversed: even without dispersing, a resident subordinate male may eventually reproduce with successive dominant females as they turn over. Thus, sons should benefit more from philopatry than daughters, while daughters should benefit more from dispersal. Given equal dispersal costs for both sexes, this makes female-biased dispersal a viable strategy. To summarize, in the studied mole vole population, females experience higher mortality than males, paying high survival costs for both reproduction and dispersal. We propose that the cooperative breeding in combination with greater breeder turnover among females than males, drives female-biased dispersal. In this scenario, male philopatry is a precondition for, rather than a consequence of the strong female-female competition and facultative polyandry. This reversal of the typical mammalian pattern warrants further investigation through genetic analyses and long-term monitoring.

## Author statements

## Acknowledgements

We are grateful to M.M. Gribanova, A.I. Rudyk, I.A. Volodin, E.V. Volodina, A.A. Panyutina, I.I. Boyarinova, M.V. Vdovina, P.S. Cherepenko, B.A. Belov, and E.D. Kolosova for their participation in our long-term field study. We thank A.V. Tchabovsky and V.A. Lukhtanov for their great help in the organization of field work. Special thanks to Andreas Spiess for his well-documented tutorial on microelectronic craft.

## Author contributions

Conceptualization: A.E.N., A.O.F., A.V.S; sample collection: A.E.N., V.R.N., A.M.B., M.M.D, A.O.F., and A.V.S.; methodology: A.E.N., A.O.F., A.M.B., and A.V.S.; investigation: A.E.N., A.O.F., A.M.B., and A.V.S.; funding acquisition: A.V.S.; writing – original draft: A.E.N., A.O.F. and A.V.S.; writing – review and editing: A.E.N., V.R.N., A.M.B., M.M.D., A.O.F., and A.V.S.

## Ethical approval

All procedures involving animals were in compliance with the national laws of the Russian Federation. The experimental protocols used in this study were approved by the Specialized Ethics Committee for Animal Research of the St. Petersburg State University (№ 131-03-9 22 November 2021).

## Consent for publication

All authors consent to the publication of this manuscript.

## Data availability

Data generated or analyzed during this study are provided in full within the published article and its supplementary materials.

## References

Bergaliev, A. M., Naumova, A. E., Nikonova, V. R., Fedosov, A. O., Rudyk, A. I., Gribanova, M. D., Dymskaya, M. M., Romanovich, A. E., and Smorkatcheva, A. V. 2026. Within-group kinship and mating system in a cooperative breeder, the northern mole vole (*Ellobius talpinus*), in the Lower Volga region. Mamm. Biol. 10.1007/s42991-026-00597-0

Boonstra, R., Krebs, C. J., Gaines, M. S., Johnson, M. L., and Craine, I. T. M. 1987. Natal Philopatry and Breeding Systems in Voles (*Microtus* Spp.). J. Anim. Ecol. 56(2): 655–673. 10.2307/5075

Borkowska A. 2011. Seasonal variation of reproductive success under female philopatry and male-biased dispersal in a common vole population. Behav. Processes. 86(1): 39–45. 10.1016/j.beproc.2010.08.005

Bray, T. C., Bloomer, P., O’Riain, M. J., and Bennett, N. C. 2012. How attractive is the girl next door? An assessment of spatial mate acquisition and paternity in the solitary cape dune mole-rat, *Bathyergus suillus*. PLoS One. 7: e39866. 10.1371/journal.pone.0039866

Brett, R. A. 1991. The population structure of naked mole-rat colonies. In: Sherman, P. W., Jarvis, J. U. M. and Alexander R. D. (eds.), The biology of the naked mole-rat. Princeton University Press, Princeton. pp. 97–136. 10.1515/9781400887132-007

Briner, T., Favre, N., Nentwig, W., and Airoldi, J.-P. 2007. Population dynamics of *Microtus arvalis* in a weed strip. Mamm. Biol. 72(2): 106–115. 10.1016/j.mambio.2006.07.006

Brom, T., Massot, M., Legendre, S., and Laloi, D. 2016. Kin competition drives the evolution of sex-biased dispersal under monandry and polyandry, not under monogamy. Anim. Behav. 113: 157–166. 10.1016/j.anbehav.2016.01.003

Buckland, S. T., Burnham, K. P., and Augustin, N. H. 1997. Model selection: an integral part of inference. Biometrics. 53(2): 603–618. 10.2307/2533961

Burnham, K. P., and Anderson, D. R. 2002. Model selection and multimodel inference: a practical information-theoretic approach. 2nd Edition, Springer-Verlag, New York. 10.1007/b97636

Burnham, K. P., and Anderson, D. R. 2004. Multimodel Inference: understanding AIC and BIC in model selection. Sociol. Methods Res. 33(2): 261–304. 10.1177/0049124104268644

Busch, C., Antinuchi, C. D., Del Valle, J. C., Kittlein, M. J., Malizia, A. I., and Vassallo, A. I. 2000. Population ecology of subterranean rodents. In: Lacey, E. A., Patton, J. L., Cameron, G. N. (eds.), Life underground: the biology of subterranean rodents. University of Chicago Press, Chicago. pp. 183–226.

Clark, A. B. 1978. Sex ratio and local resource competition in a prosimian primate. Science. 201(4351): 163–165. 10.1126/science.201.4351.163

Cockburn, A., Scott, M. P., and Dickman, C. R. 1985. Sex ratio and intrasexual kin competition in mammals. Oecologia. 66: 427–429. 10.1007/BF00378310

Crowley, P. H. 2000. Sexual dimorphism with female demographic dominance: Age, size, and sex ratio at maturation. Ecology. 81(9): 2592–2607. 10.1890/0012-9658(2000)081[2592:SDWFDD]2.0.CO;2

Cutrera, A. P., Lacey, E. A., and Busch, C. 2005. Genetic structure in a solitary rodent (*Ctenomys talarum*): implications for kinship and dispersal. Mol. Ecol. 14: 2511–2523. 10.1111/j.1365-294X.2005.02551.x

Davis-Born, R., and Wolff, J. O. 2000. Age- and sex-specific responses of the gray-tailed vole, *Microtus canicaudus*, to connected and unconnected habitat patches. Can. J. Zool. 78(5): 864–870. 10.1139/z00-017

Donald, P. F. 2007. Adult sex ratios in wild bird populations. Ibis. 149(4): 671–692. 10.1111/j.1474-919X.2007.00724.x

Dymskaya, M. M., Volodin, I. A., Smorkatcheva, A. V., Rudyk, A. I., and Volodina, E. V. 2025. Field tests reveal acoustic variation of call types in a subterranean rodent, the northern mole vole *Ellobius talpinus*. J. Mammal. 106: 237–251. 10.1093/jmammal/gyae123

Evdokimov, N. G. 1997. Dynamics of abundance of the northern mole vole (*Ellobius talpinus* Pall.). Dokl. Biol. Sci. 356: 424–426. [in Russian]

Evdokimov, N. G. 1999. Analysis of dispersal in populations of the northern mole vole (*Ellobius talpinus* Pall.). Russian Journal of Ecology. 5: 323–328. [in Russian]

Evdokimov, N. G. 2001. Population ecology of northern mole vole (*Ellobius talpinus*). Ural Branch of RAS, Yekaterinburg. [in Russian]

Evdokimov, N. G. 2013. Structure of the mole vole (*Ellobius talpinus*, Rodentia, Cricetidae) colonies. Zool. Zh. 92(3): 325–336. [in Russian] 10.7868/S0044513413030082

Evdokimov, N. G., and Pozmogova, V. P. 1984. Comparative characteristics of three populations of the northern mole vole (*Ellobius talpinus*) (the Southern Urals, Trans-Urals, and Northern Kazakhstan). In Dobrinskii, L. N. (eds.), Population ecology and morphology of mammals. Ural Scientific Centre, Sverdlovsk. pp. 103–112. [in Russian]

Evdokimov, N. G., and Pozmogova, V. P. 1992. Mountain and champaign populations of the northern mole vole (*Ellobius talpinus*) (the Southern Urals and Trans-Urals). In: Ecology of mammals of the Ural Mountains. Ural Branch of RAS, Yekaterinburg. pp. 100–119. [in Russian]

Evdokimov, N. G., and Pozmogova, V. P. 1993. Population structure of the northern mole vole (*Ellobius talpinus* Pall.) in the Transural region. Russian Journal of Ecology. 5: 53–60.

Fernández-Stolz, G. P., Stolz, J. F. B., and de Freitas, T. R. O. 2007. Bottlenecks and dispersal in the tuco-tuco das dunas, *Ctenomys flamarioni* (Rodentia: Ctenomyidae), in southern Brazil. J. Mammal. 88(4): 935–945. 10.1644/06-MAMM-A-210R1.1

Finn, K. T. 2021. Potential use of a magnetic compass during long-distance dispersal in a subterranean rodent. J. Mammal. 102(1): 250–257. 10.1093/jmammal/gyaa163.

Finn, K. T. 2022. Sociality in african mole-rats: exploring how rainfall affects dispersal and genetic exchange in the natal mole-rat (Cryptomys hottentotus natalensis). PhD thesis. University of Pretoria, Pretoria, South Africa.

Finn, K. T., Thorley J., Bensch, H. M., and Zöttl M. 2022. Subterranean life-style does not limit long distance dispersal in african mole-rats. Front. Ecol. Evol. 10:879014. 10.3389/fevo.2022.879014

Fisher, D. O., and Blomberg, S. P. 2011. Costs of reproduction and terminal investment by females in a semelparous marsupial. PLoS ONE. 6(1): e15226. 10.1371/journal.pone.0015226

Francioli, Y., Thorley, J., Finn, K., Clutton-Brock, T., and Zöttl M. 2020. Breeders are less active foragers than non-breeders in wild damaraland mole-rats. Biol. Lett. 16(10): 20200475. 10.1098/rsbl.2020.0475

Froy, H., Walling, C. A., Pemberton, J. M., Clutton-Brock, T. H., and Kruuk, L. E. B. 2016. Relative costs of offspring sex and offspring survival in a polygynous mammal. Biol. Lett. 12(9): 20160417. 10.1098/rsbl.2016.0417

Gauffre, B., Petit, E., Brodier, S., Bretagnolle, V., and Cosson, J. F. 2009. Sex-biased dispersal patterns depend on the spatial scale in a social rodent. Proc. R. Soc. B. 276(1672): 3487–3494. 10.1098/rspb.2009.0881

Getz, L., Mcguire, B., Hofmann, J., Pizzuto, T., and Frase, B. 1994. Natal dispersal and philopatry in prairie voles (*Microtus ochrogaster*): settlement, survival, and potential reproductive success. Ethol. Ecol. Evol. 6: 267–284. 10.1080/08927014.1994.9522980

Ginther, S. C., Cameron, H., White, C. R., and Marshall, D. J. 2024. Metabolic loads and the costs of metazoan reproduction. Science. 384: 763–767. 10.1126/science.adk6772

Golov, B. A. 1954. The live-trap for the mole vole. Byull. Mosk. O-va. Ispyt. Prir. Biol. 59: 95–96 [in Russian]

Graw, B., Lindholm, A. K., and Manser, M. B. 2016. Female-biased dispersal in the solitarily foraging slender mongoose, *Galerella sanguinea*, in the Kalahari. Anim. Behav. 111: 69–78. 10.1016/j.anbehav.2015.09.026

Greenwood, P. J. 1980. Mating systems, philopatry and dispersal in birds and mammals. Anim. Behav. 28(4): 1140–1162. 10.1016/s0003-3472(80)80103-5

Gros, A., Poethke, H. J., and Hovestadt, T. 2009. Sex-specific spatio-temporal variability in reproductive success promotes the evolution of sex-biased dispersal. Theor. Popul. Biol. 76(1): 13–18. 10.1016/j.tpb.2009.03.002

Grüebler, M. U., and Naef-Daenzer, B. 2008. Postfledging parental effort in barn swallows: evidence for a trade-off in the allocation of time between broods. Anim. Behav. 75(6): 1877–1884. 10.1016/j.anbehav.2007.12.002

Hammond, R. L., Handley, L. J. L., Winney, B. J., Bruford, M. W., and Perrin, N. 2006. Genetic evidence for female-biased dispersal and gene flow in a polygynous primate. Proc. R. Soc. B. 273(1585): 479–484. 10.1098/rspb.2005.3257

Hazell, R. W. A., Bennett, N. C., Jarvis, J. U. M., and Griffin, M. 2000. Adult dispersal in the co-operatively breeding damaraland mole-rat (*Cryptomys damarensis*): a case study from the Waterberg region of Namibia. J. Zool. 252(1): 19–25. 10.1111/j.1469-7998.2000.tb00816.x

Heske, E. J. 1987. Spatial structuring and dispersal in a high density population of the california vole *Microtus californicus*. Ecography. 10(2): 137–145. 10.1111/j.1600-0587.1987.tb00750.x

Houslay, T. M., Vullioud, P., Zöttl, M., and Clutton-Brock, T. H. 2020. Benefits of cooperation in captive damaraland mole-rats. Behav. Ecol. 31(3): 711–718. 10.1093/beheco/araa015

Jannett, F. J. 1981. Sex ratios in high-density populations of the montane vole, *Microtus montanus*, and the behavior of territorial males. Behav. Ecol. Sociobiol. 8: 297–307. 10.1007/BF00299530

Johnson, M. L., and Gaines, M. S. 1990. Evolution of dispersal: theoretical models and empirical tests using birds and mammals. Annu. Rev. Ecol. Syst. 21(1): 449–480. 10.1146/annurev.es.21.110190.002313

Jones, O. R., and Wang, J. 2010. COLONY: a program for parentage and sibship inference from multilocus genotype data. Mol. Ecol. Resour. 10(3): 551–555. 10.1111/j.1755-0998.2009.02787.x

Kawalika, M., and Burda, H. 2007. Giant mole-rats, *Fukomys mechowii*, 13 years on the stage. In: Begall, S., Burda, H., Schleich, C. E. (eds.), Subterranean Rodents. Springer, Berlin. pp. 205–219. 10.1007/978-3-540-69276-8_15

Kryštufek, B., and Shenbrot, G. 2022. Voles and lemmings (Arvicolinae) of the palaearctic region. University of Maribor Press, Maribor. 10.18690/um.fnm.2.2022

Kuprina, K. V., and Smorkatcheva, A. V. 2019. Noninvasive age estimation in rodents by measuring incisors width, with the zaisan mole vole (*Ellobius tancrei*) as an example. Mammalia. 83(1): 64–69. 10.1515/mammalia-2017-0163

Laake, J. L. 2013. RMark: an R interface for analysis of capture–recapture data with MARK. AFSC Processed Report 2013-01. Alaska Fisheries Science Center, National Marine Fisheries Service, NOAA, Seattle, WA. 25 p. 10.32614/CRAN.package.RMark

Lacey, E. A., and Wieczorek, J. R. 2004. Kinship in colonial tuco-tucos: Evidence from group composition and population structure. Behav. Ecol. 15(6): 988–996. 10.1093/beheco/arh104

Lambin, X. 1994. Natal philopatry, competition for resources, and inbreeding avoidance in townsend’s voles (*Microtus townsendii*). Ecology. 75(1): 224–235. 10.2307/1939396

Le Galliard, J.-F., Rémy, A., Ims, R. A., and Lambin, X. 2012. Patterns and processes of dispersal behaviour in arvicoline rodents. Mol. Ecol., 21(3): 505–523. 10.1111/j.1365-294X.2011.05410.x

Lebreton, J.-D., and Pradel, R. 2002. Multistate recapture models: modelling incomplete individual histories. J. Appl. Stat., 29(1–4): 353–369. 10.1080/02664760120108638

Lebreton, J.-D., Burnham, K., Clobert, J., and Anderson, D. R. 1992. Modeling survival and testing biological hypotheses using marked animals: a unified approach with case studies. Ecol. Monogr. 62: 67–118. 10.2307/2937171

Lehmann, L., and Perrin, N. 2003. Inbreeding avoidance through kin recognition: choosy females boost male dispersal. Am. Nat., 162(5): 638–652. 10.1086/378823

Liker, A., and Székely, T. 2005. Mortality costs of sexual selection and parental care in natural populations of birds. Evolution. 59(4): 890–897. 10.1111/j.0014-3820.2005.tb01762.x

Liker, A., Freckleton, R. P., Remeš, V., and Székely, T. 2015. Sex differences in parental care: gametic investment, sexual selection, and social environment. Evolution. 69(11): 2862–2875. 10.1111/evo.12786

Maag, N., Paniw, M., Cozzi, G., Manser, M., Clutton-Brock, T. H., and Ozgul, A. 2022. Dispersal decreases survival but increases reproductive opportunities for subordinates in a cooperative breeder. Am. Nat. 199(5): 679–690. 10.1086/719029

Mabry, K. E., Shelley, E. L., Davis, K. E., Blumstein, D. T., and Van Vuren, D. H. 2013. Social mating system and sex-biased dispersal in mammals and birds: a phylogenetic analysis. PLoS ONE. 8(3): e57980. 10.1371/journal.pone.0057980

Małek, D. K., Dańko, M. J., and Czarnołęski, M. 2023. Effect of age, mating history and temperature on male reproductive costs in the bean beetle *Callosobruchus maculatus*. J. Stored Prod. Res. 102: 102110. 10.1016/j.jspr.2023.102110

McHuron, E. A., Adamczak, S., Costa, D. P., and Booth, C. 2023. Estimating reproductive costs in marine mammal bioenergetic models: a review of current knowledge and data availability. Conserv. Physiol. 11(1): coac080. 10.1093/conphys/coac080

Metcalf, C. J. E., and Graham, A. L. 2018. Schedule and magnitude of reproductive investment under immune trade-offs explains sex differences in immunity. Nat. Commun. 9: 4391. 10.1038/s41467-018-06793-y

Miranda, O. G., Colchero, F., Valdebenito, J. O., Cortez, D., Conde, D. A., Pipoly, I., Liker, A., Vági, B., Bertelsen, M. F., Kilili, A., Urrutia, A. O., and Székely, T. 2025. Biased birth sex ratios of mammals and birds in zoos. Sci. Rep. 15(1): 20506. 10.1038/s41598-025-05039-4

Mora, M. S., Mapelli, F. J., Gaggiotti, O. E., Kittlein, M. J., Lessa, E. P. 2010. Dispersal and population structure at different spatial scales in the subterranean rodent *Ctenomys australis*. BMC Genet. 11: 9. 10.1186/1471-2156-11-9

Myers, J. H., and Krebs, C. J. 1971. Sex ratios in open and enclosed vole populations: Demographic implications. Am. Nat. 105(944): 325–344. 10.1086/282728

Mynhardt, S., Harris-Barnes, L., Bloomer, P., and Bennett, N. C. 2021. Spatial population genetic structure and colony dynamics in Damaraland mole-rats (*Fukomys damarensis*) from the southern Kalahari. BMC Ecol Evol. 21: 221. 10.1186/s12862-021-01950-2

Nikonova, V. R., Naumova, A. E., Bergaliev, A. M., Dymskaya, M. M., Rudyk, A. I., Volodina, E. V., and Smorkatcheva, A. V. 2024. Dental radiography as a low-invasive field technique to estimate age in small rodents, with the mole voles (*Ellobius*) as an example. Eur. J. Wildl. Res. 70: 46. 10.1007/s10344-024-01802-6

Novikov, E. A. 2007. Frugal strategy as a base of mole-vole (*Ellobius talpinus*: Rodentia) adaptations to the fossil war of life. Zh. Obshch. Biol. 68(4): 267–274. [in Russian]

Novikov, E. A., Petrovskii, D. V., and Moshkin, M. P. 2007. Population structure of the northern mole vole at the northeastern periphery of its distribution range. Sib. Ecol. Zh. 4: 669–676. [in Russian]

O’Riain, M. J., Jarvis, J. U. M., and Faulkes, C. G. 1996. A dispersive morph in the naked mole-rat. Nature. 380: 619–621. 10.1038/380619a0

Patzenhauerová, H., Bryja, J., and Šumbera, R. 2010. Kinship structure and mating system in a solitary subterranean rodent, the silvery mole-rat. Behav. Ecol. Sociobiol. 64: 757–767. 10.1007/s00265-009-0893-4

Patzenhauerová, H., Skilba, J., Bryja, J., and Šumbera, R. 2013. Parentage analysis of Ansell’s mole-rat family groups indicates a high reproductive skew despite relatively relaxed ecological constraints on dispersal. Mol. Ecol. 22(19): 4988–5000. 10.1111/mec.12434

Payevsky, V. A. 2020. Sex structure and sex-specific survival in bird populations (Review). Zh. Obshch. Biol. 81(4): 272–284. [in Russian]

Payevsky, V. A. 2021. Sex ratio and sex-specific survival in avian populations: a review. Biol. Bull. Rev. 11: 317–327. 10.1134/S2079086421030099

Perrin, N., and Mazalov, V. 1999. Dispersal and inbreeding avoidance. Am. Nat. 154(3): 282–292. 10.1086/303236

Perrin, N., and Mazalov, V. 2000. Local competition, inbreeding, and the evolution of sex-biased dispersal. Am. Nat. 155(1): 116–127. 10.1086/303296

Pickett, E. J., Stockwell, M. P., Pollard, C. J., Garnham, J. I., Clulow, J., and Mahony, M. J. 2012. Estimates of sex ratio require the incorporation of unequal catchability between sexes. Wildl. Res. 39(4): 350–354. 10.1071/wr11193

Pipoly, I., Bókony, V., Kirkpatrick, M., Donald, P. F., Székely, T., and Liker, A. 2015. The genetic sex-determination system predicts adult sex ratios in tetrapods. Nature. 527: 91–94. 10.1038/nature15380

Promislow, D. E. L. 1992. Costs of sexual selection in natural populations of mammals. Proc. R. Soc. Lond., B. 247(1320): i–v. 10.1098/rspb.1992.0030

Puzachenko, A. Y. 1998. Demography of dispersion in mole rat *Spalax microphthalmus* (Rodentia, Spalacidae). Zool. Zh. 77(3): 364–368. [in Russian]

Romano, A., Basile, M., and Costa, A. 2018. Skewed sex ratio in a forest salamander: artefact of the different capture probabilities between sexes or actual ecological trait? Amphibia-Reptilia. 39(1): 79–86. 10.1163/15685381-17000029

Romano, A., Liker, A., Bazzi, G., Ambrosini, R., Møller, A. P., and Rubolini, D. 2022. Annual egg productivity predicts female-biased mortality in avian species. Evolution. 76(11): 2553–2565. 10.1111/evo.14623

Roy, J., Gray, M., Stoinski, T., Robbins, M. M., and Vigilant, L. 2014. Fine-scale genetic structure analyses suggest further male than female dispersal in mountain gorillas. BMC Ecol. 14: 21. 10.1186/1472-6785-14-21

Rudyk, A. I., Kuprina, K., Bergaliev, A. M., Galkina, S. A., Romanovich, A. E., Novikov, E. A., Volodina, E. V., and Smorkatcheva, A. V. 2025. Genetic diversity and population structure of the subterranean rodent, northern mole vole (*Ellobius talpinus*). Mamm. Biol. 105: 571–588. 10.1007/s42991-025-00498-8

Salvioni, M., and Lidicker, W. Z. Jr. 1995. Social organization and space use in California voles: seasonal, sexual, and age-specific strategies. Oecologia. 101(4): 426–438. 10.1007/BF00329421

Sandercock, B. K. 2006. Estimation of demographic parameters from live-encounter data: a summary review. J. Wildl. Manage. 70(6): 1504–1520. 10.2193/0022-541x(2006)70[1504:eodpfl]2.0.co;2

Sharp, S. P., and Clutton-Brock, T. H. 2011. Competition, breeding success and ageing rates in female meerkats. J. Evol. Biol. 24(8): 1756–1762. 10.1111/j.1420-9101.2011.02304.x

Shevlyuk, N. N., and Yelina, E. E. 2008. Biology of reproduction of the northern mole vole Ellobius talpinus. Orenburg State Pedagogical University Press, Orenburg. [in Russian]

Shubin, I. G. 1961. On the ecology of the northern mole vole in central Kazakhstan. Zool. Zh. 40(10): 1543–1551. [in Russian]

Smorkatcheva, A. V. 2003. Parental care in the captive mandarin vole, *Lasiopodomys mandarinus*. Can. J. Zool. 81(8): 1339–1348. 10.1139/z03-100

Smorkatcheva, A. V., and Kumaitova, A. R. 2013. Delayed dispersal in the zaisan mole vole (*Ellobius tancrei*): helping or extended parental investment? J. Ethol. 32: 53–61. 10.1007/s10164-013-0392-y

Smorkatcheva, A. V., and Kuprina, K. 2018. Does sire replacement trigger plural reproduction in matrifilial groups of a singular breeder, *Ellobius tancrei*? Mamm. Biol. 88: 144–150. 10.1016/j.mambio.2017.09.005

Speakman, J. R. 2008. The physiological costs of reproduction in small mammals. Philos. Trans. R. Soc. Lond., B. 363(1490): 375–398. 10.1098/rstb.2007.2145

Staerk, J., Conde, D. A., Tidière, M., Lemaître, J. F., Liker, A., Vági, B., Pavard, S., Giraudeau, M., Smeele, S. Q., Vincze, O., Ronget, V., da Silva, R., Pereboom, Z., Bertelsen, M. F., Gaillard, J. M., Székely, T., and Colchero, F. 2025. Sexual selection drives sex difference in adult life expectancy across mammals and birds. Sci. Adv. 11(40): eady8433. 10.1126/sciadv.ady8433

Šumbera, R., Šklíba, J., Elichová, M., Chitaukali, W. N., and Burda, H. 2007. Natural history and burrow system architecture of the silvery mole-rat from Brachystegia woodland. J. Zool. 274(1): 77–84. 10.1111/j.1469-7998.2007.00359.x

Sweitzer, R. A., and Berger, J. 1998. Evidence for female-biased dispersal in North American porcupines (*Erethizon dorsatum*). J. Zool. 244: 159–166. 10.1017/s0952836998002015

Székely, T., Weissing, F. J., and Komdeur, J. 2014. Adult sex ratio variation: implications for breeding system evolution. J. Evol. Biol. 27(8): 1500–1512. 10.1111/jeb.12415

Tamarin, R. H. 1977. Demography of the beach vole (*Microtus breweri*) and the meadow vole (*M. pennsylvanicus*) in southeastern Massachusetts. Ecology. 58(6): 1310–1321. 10.2307/1935083

Vasjukov, V. M., Senator, S. A., Saksonov, S. V., Zibzeev, E. G., and Korolyuk, A. Yu. 2018. Materials to the flora of natural monument “Diakovskiy Forest” (Saratov Region). Bull. Bot. Gard. Saratov State Univ. 16(3): 3–18. [in Russian]

Végvári, Z., Katona, G., Vági, B., Freckleton, R. P., Gaillard, J. M., Székely, T., and Liker, A. 2018. Sex-biased breeding dispersal is predicted by social environment in birds. Ecol. Evol. 8(13): 6483–6491. 10.1002/ece3.4095

Vleck, D. 1979. The energy cost of burrowing by the pocket gopher *Thomomys bottae*. Physiol. Zool. 52(2): 122– 136. 10.1086/physzool.52.2.30152558

Waser, P. M., Austad, S. N., and Keane, B. 1986. When should animals tolerate inbreeding? Am. Nat. 128(4): 529–537. 10.1086/284585

Welbergen, J. A. 2010. Growth, bimaturation, and sexual size dimorphism in wild gray-headed flying foxes (*Pteropus poliocephalus*). J. Mammal. 91(1): 38–47. 10.1644/09-MAMM-A-157R.1

White, G. C., and Burnham, K. P. 1999. Program MARK: Survival estimation from populations of marked animals. Bird Study. 46(1): S120–S139. 10.1080/00063659909477239

Williams, G. C. 1966. Natural selection, the costs of reproduction, and a refinement of Lack’s principle. Am. Nat. 100: 687– 690. 10.1086/282461

Winder, L. A., Simons, M. J. P., and Burke, T. 2025. No evidence for a trade-off between reproduction and survival in a meta-analysis across birds. eLife. 12: RP87018. 10.7554/eLife.87018.5

Zenuto, R. R., and Busch, C. 1998. Population biology of the subterranean rodent *Ctenomys australis* (Tuco-tuco) in a coastal dunefield in Argentina. Z. Säugetierkunde. 63: 357–367. https://www.biodiversitylibrary.org/part/192359

Zöttl, M., Bensch, H. M., Finn, K. T., Hart, D. W., Thorley, J., Bennett, N. C., and Braude, S. 2022. Capture order across social Bathyergids indicates similarities in division of labour and spatial organisation. Front. Ecol. Evol. 10: 877221. 10.3389/fevo.2022.877221

