## Supplemental file for "Apparent survival suggests the female-biased dispersal in a truly subterranean rodent, *Ellobius talpinus*"

**Table S1** Detailed numerical annual survival rates and 95% confidence intervals for each demographic group and each period based on model averaging. F – female, M – male, Y – young-of-the-year, B – breeder, N – non-breeder.

| Sex | Stratum | Time period | $\varphi$ estimate | se | 95% CI |
|---|---|---|---|---|---|
| F | Y | 2022 - 2023 | 0.19 | 0.05 | 0.12 - 0.30 |
| F | B | 2022 - 2023 | 0.33 | 0.09 | 0.18 - 0.52 |
| F | N | 2022 - 2023 | 0.32 | 0.09 | 0.18 - 0.52 |
| M | Y | 2022 - 2023 | 0.34 | 0.06 | 0.24 - 0.47 |
| M | B | 2022 - 2023 | 0.52 | 0.09 | 0.34 - 0.69 |
| M | N | 2022 - 2023 | 0.50 | 0.08 | 0.35 - 0.66 |
| F | Y | 2023 - 2024 | 0.18 | 0.04 | 0.12 - 0.27 |
| F | B | 2023 - 2024 | 0.31 | 0.08 | 0.18 - 0.48 |
| F | N | 2023 - 2024 | 0.31 | 0.08 | 0.18 - 0.48 |
| M | Y | 2023 - 2024 | 0.33 | 0.05 | 0.24 - 0.43 |
| M | B | 2023 - 2024 | 0.50 | 0.09 | 0.34 - 0.66 |
| M | N | 2023 - 2024 | 0.48 | 0.07 | 0.36 - 0.61 |
| F | Y | 2024 - 2025 | 0.17 | 0.04 | 0.11 - 0.26 |
| F | B | 2024 - 2025 | 0.30 | 0.08 | 0.18 - 0.47 |
| F | N | 2024 - 2025 | 0.30 | 0.08 | 0.17 - 0.47 |
| M | Y | 2024 - 2025 | 0.32 | 0.05 | 0.23 - 0.42 |
| M | B | 2024 - 2025 | 0.48 | 0.08 | 0.33 - 0.65 |
| M | N | 2024 - 2025 | 0.47 | 0.07 | 0.35 - 0.60 |

**Table S2** Detailed numerical transition probabilities and 95% confidence intervals for each demographic group based on model averaging. F – female, M – male, Y – young-of-the-year, B – breeder, N – non-breeder.

| Sex | From | To | $\psi$ estimate | se | 95% CI |
|---|---|---|---|---|---|
| F | Y | B | 0.29 | 0.11 | 0.13 - 0.54 |
| F | Y | N | 0.71 | 0.11 | 0.46 - 0.87 |
| F | B | B | 1.00 | 0.00 | 1.00 - 1.00 |
| F | B | N | 0.00 | 0.00 | 0.00 - 0.00 |
| F | N | B | 0.43 | 0.19 | 0.14 - 0.77 |
| F | N | N | 0.57 | 0.19 | 0.23 - 0.86 |
| M | Y | B | 0.12 | 0.07 | 0.04 - 0.32 |
| M | Y | N | 0.88 | 0.07 | 0.68 - 0.96 |
| M | B | B | 0.71 | 0.17 | 0.33 - 0.93 |
| M | B | N | 0.29 | 0.17 | 0.07 - 0.67 |
| M | N | B | 0.27 | 0.11 | 0.10 - 0.53 |
| M | N | N | 0.73 | 0.11 | 0.47 - 0.90 |

**Table S3** Input dataset for RMark analysis. Y – young-of-the-year, B – breeder, N – non-breeder.

| Encounter history | Amount: Female | Amount: Male |
|---|---|---|
| 00B0 | 0 | 1 |
| 00B0 | 5 | 0 |
| 00BB | 3 | 0 |
| 00BU | 0 | 4 |
| 00Y0 | 0 | 40 |
| 00Y0 | 37 | 0 |
| 00YB | 2 | 0 |
| 00YN | 5 | 0 |
| 00YU | 0 | 16 |
| 00N0 | 0 | 5 |
| 00N0 | 1 | 0 |
| 00NB | 1 | 0 |
| 00NU | 0 | 5 |
| 0B00 | 0 | 1 |
| 0B00 | 1 | 0 |
| 0BB0 | 0 | 3 |
| 0BB0 | 2 | 0 |
| 0Y00 | 0 | 27 |
| 0Y00 | 31 | 0 |
| 0YB0 | 0 | 2 |
| 0YB0 | 2 | 0 |
| 0YBU | 0 | 1 |
| 0YN0 | 0 | 7 |
| 0YN0 | 1 | 0 |
| 0YNB | 1 | 0 |
| 0YNN | 1 | 0 |
| 0YNU | 0 | 4 |
| 0N00 | 0 | 4 |
| 0N00 | 6 | 0 |
| 0NB0 | 1 | 0 |
| 0NN0 | 0 | 2 |
| B000 | 0 | 4 |
| B000 | 2 | 0 |
| BB00 | 0 | 1 |
| BB00 | 3 | 0 |
| BBNU | 0 | 1 |
| BN00 | 0 | 1 |
| Y000 | 0 | 14 |
| Y000 | 16 | 0 |
| YB00 | 1 | 0 |
| YN00 | 0 | 2 |
| YN00 | 3 | 0 |
| YNB0 | 0 | 2 |

| | | |
|---|---|---|
| YNBU | 0 | 1 |
| YNN0 | 0 | 3 |
| YNNN | 1 | 0 |
| YNNU | 0 | 2 |
| N000 | 0 | 4 |
| N000 | 1 | 0 |
| NN00 | 0 | 1 |
| NN00 | 1 | 0 |
| NNBU | 0 | 1 |
| NNN0 | 0 | 1 |

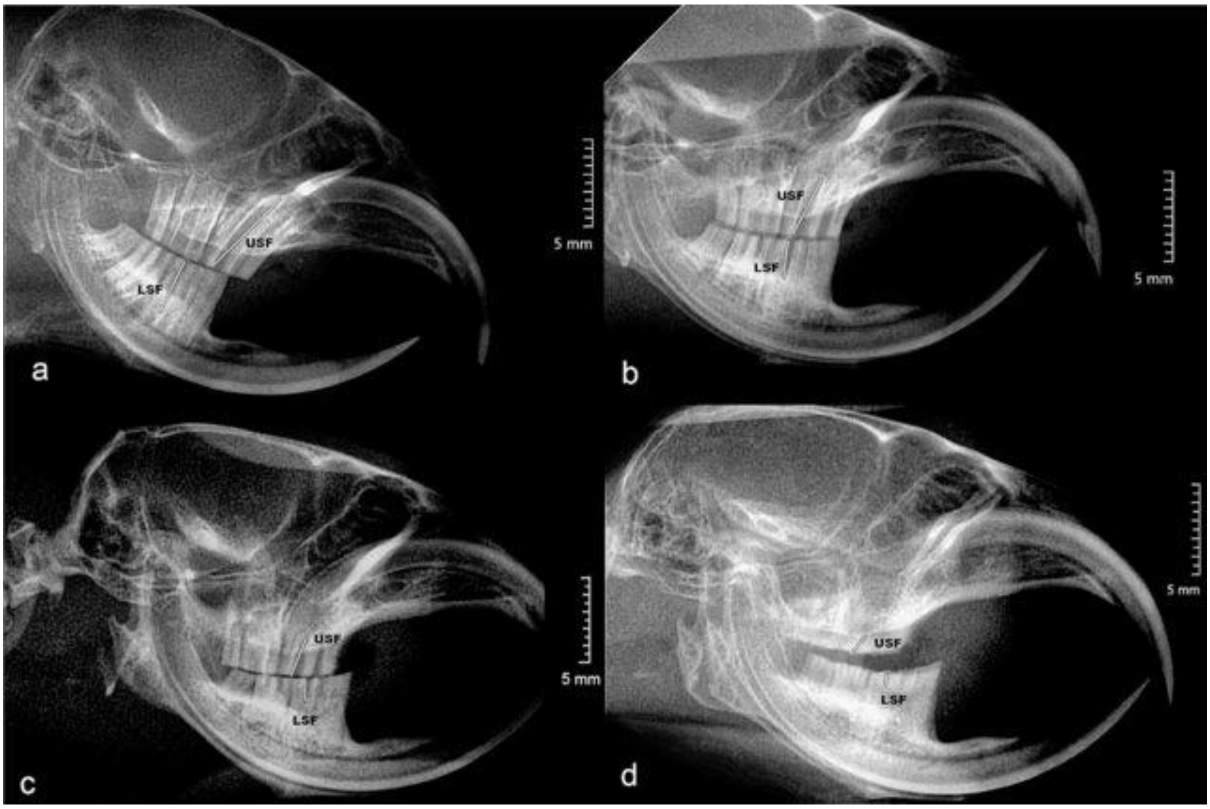


**Fig. S1** Skull radiographs of mole voles from three age classes: a - class 1 (young-of-the-year), b - class 2 (yearling); c, d - class 3 (survived two or more winters). USF – synclinal fold of the 1st upper molar; LSF – synclinal fold of the 1st lower molar from Nikonova et. al. (2024)

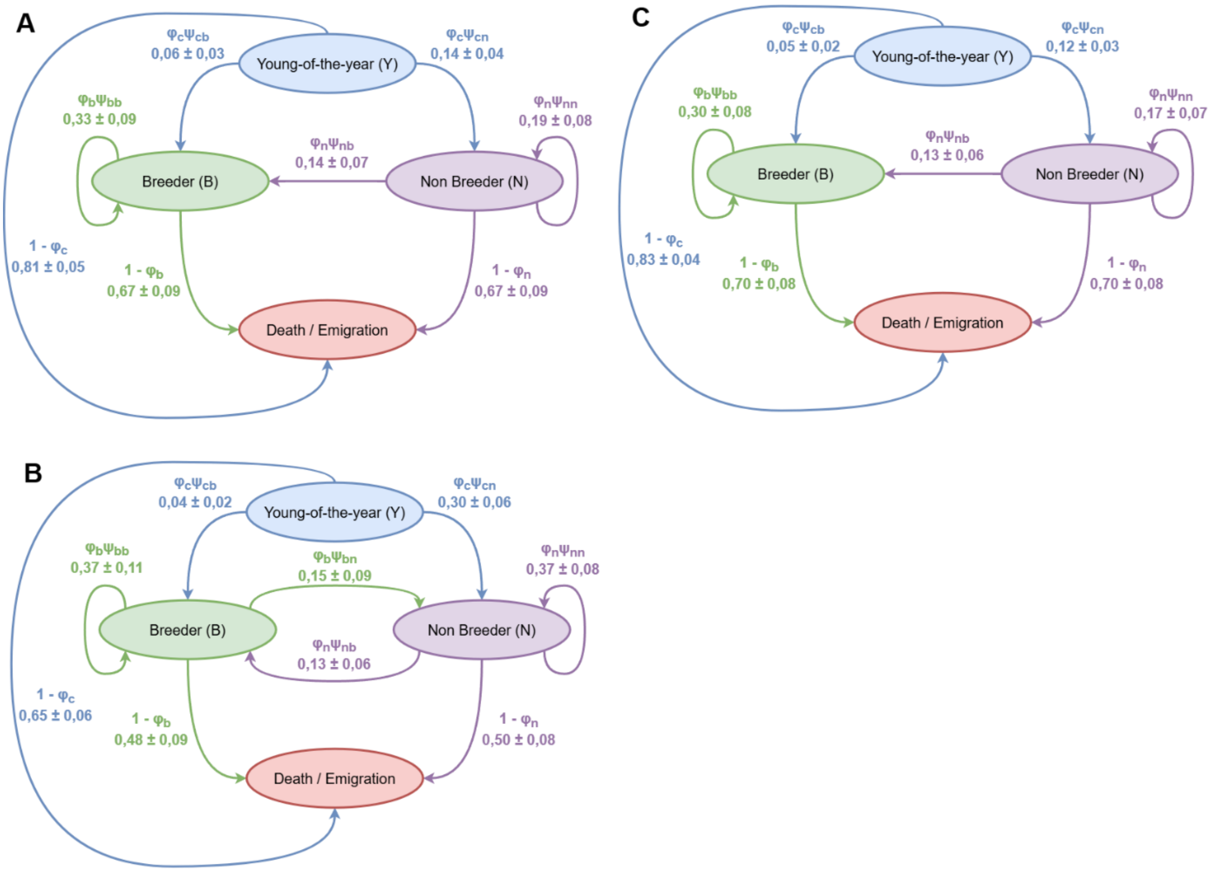


**Fig. S2 A, B, C** Graphical representation of apparent survival and transition probabilities combined for A: females (n = 28) in time period between 2022 and 2023, B: males (n = 38) in time period between 2022 and 2023, C: females (n = 58) in time period between 2024 and 2025 based on model averaging. Numbers are settled as value ± SE.

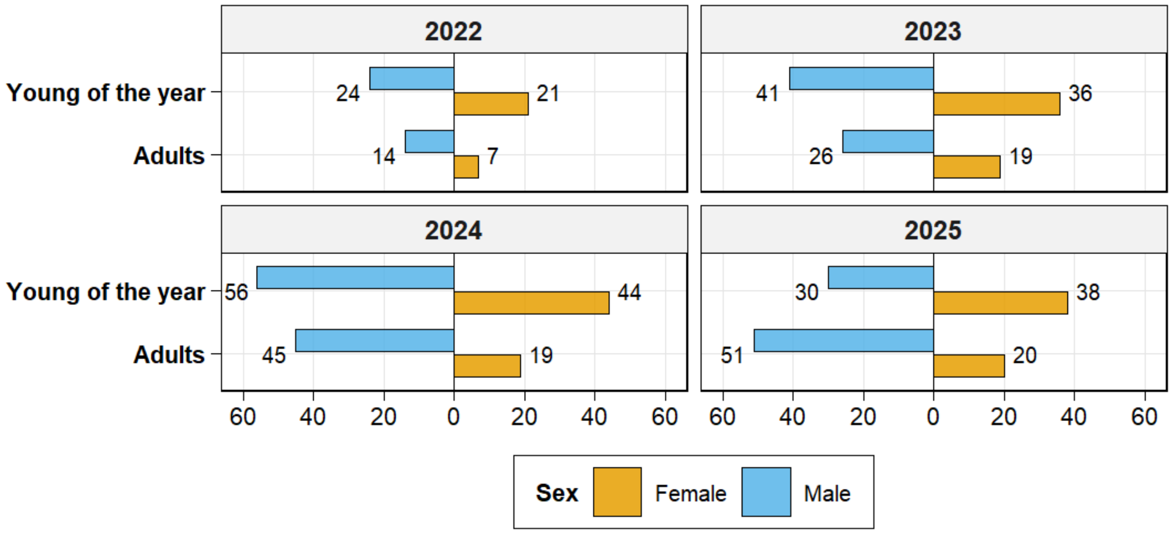


**Fig. S3** Sex ratio of young-of-the-year and adult mole voles in the focal population per year
